# Rediscovery of red wolf ghost alleles in a canid population along the American Gulf Coast

**DOI:** 10.1101/420356

**Authors:** Elizabeth Heppenheimer, Kristin E. Brzeski, Ron Wooten, Will Waddell, Linda Y. Rutledge, Michael J. Chamberlain, Daniel R. Stahler, Joseph W. Hinton, Bridgett M. vonHoldt

## Abstract

Rediscovering species once thought to be extinct or on the edge of extinction is rare. Red wolves have been extinct along the Gulf Coast region since 1980, with their last populations found in coastal Louisiana and Texas. We report the rediscovery of red wolf ghost alleles in a canid population on Galveston Island, Texas. We analyzed over 7,000 SNPs in 60 canid representatives from all legally recognized North American *Canis* species and two phenotypically ambiguous canids from Galveston Island. We found notably high Bayesian cluster assignments of the Galveston canids to captive red wolves with extensive sharing of red wolf private alleles. Today, the only known extant wild red wolves persist in a reintroduced population in North Carolina, which is dwindling amongst political and taxonomic controversy. Our rediscovery of red wolf ancestry after almost 40 years introduces both positive opportunities for additional conservation action and difficult policy challenges.

## Introduction

Red wolves (*Canis rufus*) once roamed across the southeastern United States but were declared extinct in the wild by 1980 due to habitat loss, predator control programs, disease, and interbreeding with encroaching coyotes (*Canis latrans*). In 1967, the U.S. Fish & Wildlife Service (USFWS) listed red wolves as endangered under the U.S. Endangered Species Preservation Act due to their rapid population decline in the American south, and subsequently were among the first species listed on the 1973 Endangered Species Act (ESA), the landmark of the Unites States’ environmental law (*1*). On the brink of extinction, red wolf recovery was initiated through the trapping of what was believed to be the last wild red wolves that lived along the Gulf Coast of Louisiana and Texas in the 1970s (*1-5*). Individuals were selected as founders for the captive breeding program based on morphology and behavioral traits considered to be species informative (*6,7*). Over 240 canids were trapped from coastal Louisiana and Texas between 1973 and 1977 (*6*). Forty individuals were selected for captive breeding, of which 17 were deemed 100% wolf. However, only 14 wolves successfully produced litters, which comprised the first generation of captive wolves from which all red wolves in the recovery program descend.

Due to the highly successful captive breeding program, red wolves were restored to the landscape in North Carolina less than a decade after becoming extinct in the wild (*6*). This historic event represented the first attempt to reintroduce a wild-extinct species in the United States and set a precedent for returning wild-extinct wildlife to the landscape. The success of the red wolf recovery program was the foundation upon which other wolf introductions were guided, including the gray wolf (*C. lupus*) reintroduction to the northern Rocky Mountains in Yellowstone National Park, Wyoming, and central Idaho, and the ongoing restoration efforts for the Mexican wolves (*C. lupus baileyi*) in the southwest (*8,9*). Although successful by many measures (*7*), the North Carolina experimental population (NCEP) of red wolves was reduced by the USFWS in response to negative political pressure from the North Carolina Wildlife Resource Commission and a minority of private landowners (*10*). Further, gunshot-related mortalities have increased the probability that wolf packs deteriorate before the breeding season, which facilitates the establishment of coyote-wolf breeding pairs (*11,12*). Consequently, the NCEP has fewer than 40 surviving members (*13*) and are once again on the brink of extinction in the wild.

Interbreeding between red wolves and coyotes is well documented and is viewed as a threat to red wolf recovery (*14*). When historic populations of red wolves along the Gulf Coast were surveyed, it was feared that these coastal populations were the last remnants of pre-recovery wild wolves and were likely to quickly become genetically extinct through introgressive swamping of coyote genetics (*15*). Yet, there continued to be reports of red wolves in rural coastal areas of Louisiana and Texas since the 1970s where coyotes were not considered part of the local fauna (*5,16*). Previous efforts to detect surviving red wolves or their hybrids in the region proved unsuccessful (*17*). However, the possibility remains that individuals with substantial red wolf ancestry have naturally persisted in isolated areas of the Gulf Coast. For example, body measurements of coyote-like canids in southwestern Louisiana were similar to those of confirmed red wolf-coyote hybrids in the NCEP (*18*). These individuals would harbor ghost alleles of the original red wolves, with these alleles likely lost in the contemporary red wolf population during the extreme population bottleneck, drift, and inbreeding.

For red wolf ghost alleles to persist, a remnant Gulf Coast population would need to be relatively isolated from frequent interbreeding with coyotes (*14*). Although red wolves that co-occur with coyotes in the NCEP exhibit assortative mating patterns (*19*), an island would promote low rates of genetic exchange by providing a geographic barrier and could be an ideal location for which red wolf ghost alleles would persist under limited hybridization. We report evidence that Galveston Island, TX may represent one such location. The ancestral population from which all contemporary red wolves descend were trapped from Jefferson, Chambers, southern Orange, and eastern Galveston counties in Texas and Cameron and southern Calcasieu parishes in Louisiana (*16*) (Fig. 1). Given Galveston Island’s location and isolation from the mainland, it is a probable region to harbor red wolf ghost alleles. Recent images captured of Galveston Island canids (Fig. 2) piqued interest of local naturalists and two genetic samples were taken from roadkill individuals. We conducted genomic analyses and found evidence of red wolf ancestry in modern day Galveston Island canids.

**Figure 1.**
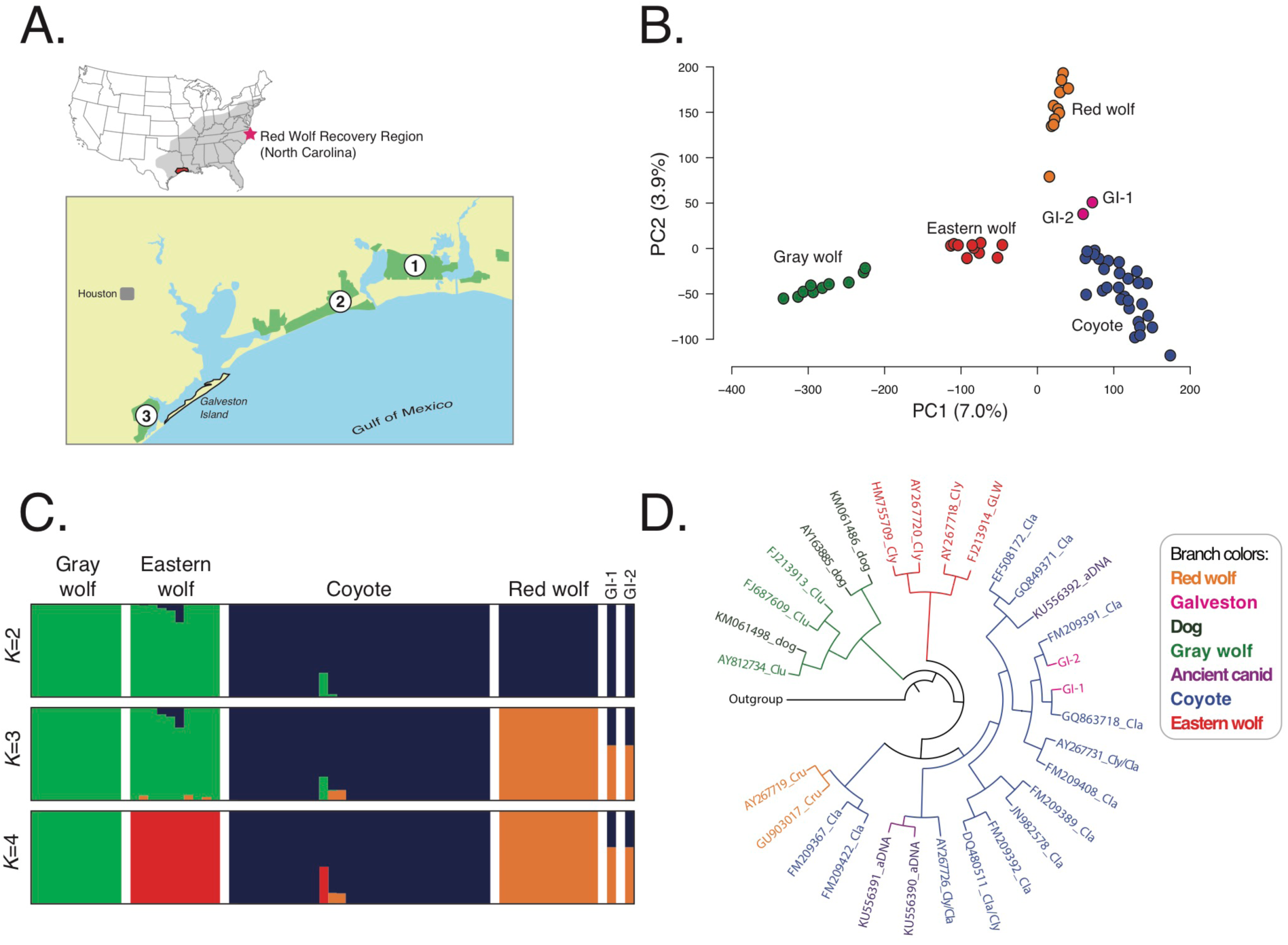
Analyses of genome wide SNP and mtDNA data across all legally recognized wild *Canis* species and two canids from Galveston Island. A) Map of area. Site 1 is Sabine National Wildlife Refuge, site 2 is McFadden National Wildlife Refuge, and site 3 is Brazoria National Wildlife Refuge. Cluster patterns were assessed across 7,068 SNPs with a B) principal component analysis and C) admixture analysis of *K*=2-4 partitions. D) Clade membership was determined by reconstruction of the Bayesian haplotype tree with the highest posterior probability (Prob=0.98) from 234bp of mtDNA sequence data from the control region; taxonomic designation of eastern wolf is based on assigned clade and sample location, not necessarily field identification.

**Figure 2.**
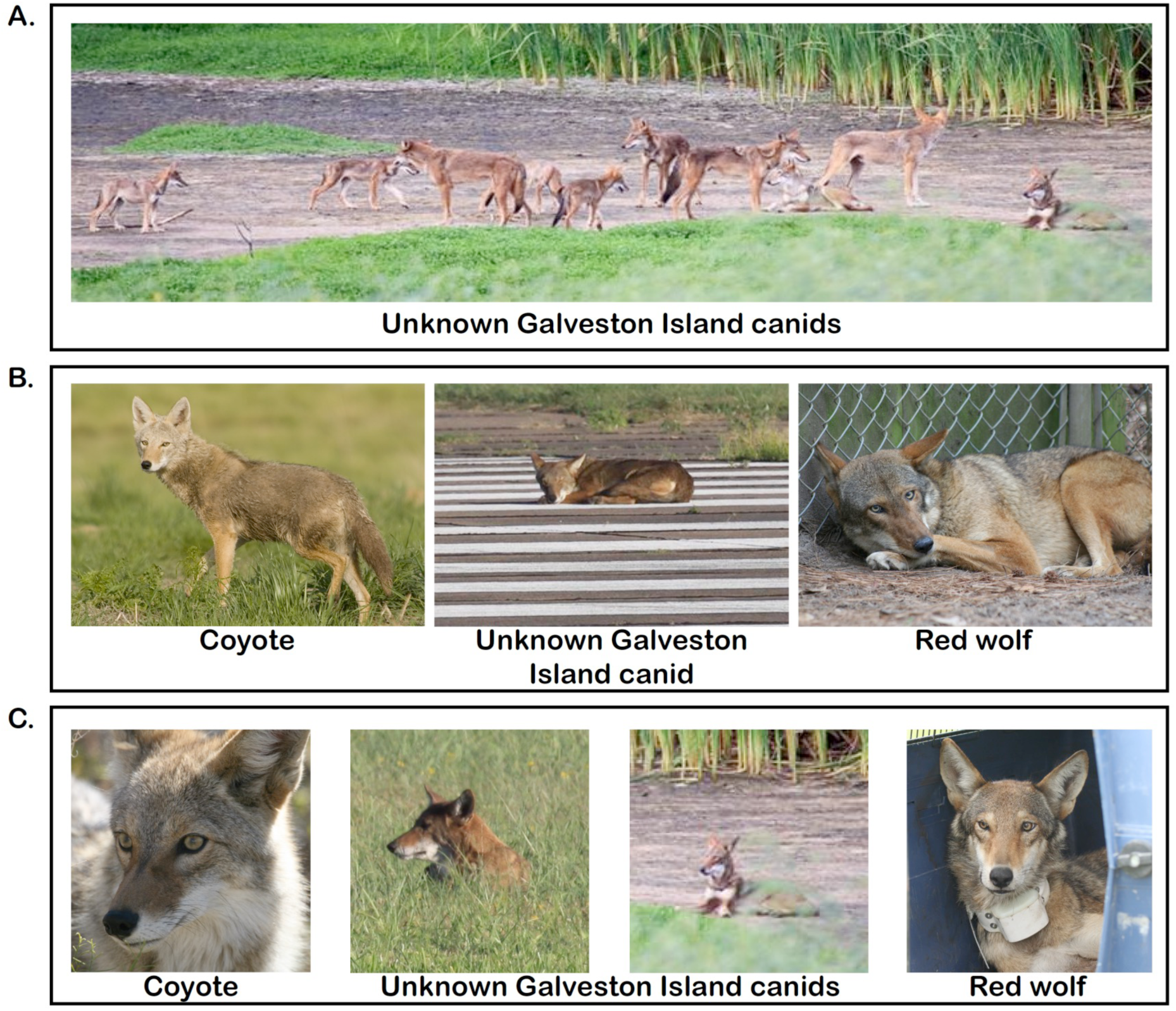
Photographic comparison of coyotes, Galveston Island (GI) canids, and red wolves. Photo credit and location as follows: A) Pack of GI canids, Galveston Island, TX, credit: R. Wooten. B) Western coyote, Intermountain West, United States, credit: Wikimedia commons, Rich Keen/DPRA. GI canid laying on airport runway, Galveston Island, TX, credit: R. Wooten. Captive female red wolf, Alligator River National Wildlife Refuge, NC, credit: R. Nordsven, USFWS. C) Western coyote, Joshua Tree National Park, CA, credit: Wikimedia commons, Michael Vamstad/NPS. Head-shots of GI canids, Galveston Island, TX, credit: R. Wooten. Wild juvenile male red wolf prior to release, Albemarle Peninsula, NC, credit R. Nordsven, USFWS.

**Figure 3.**
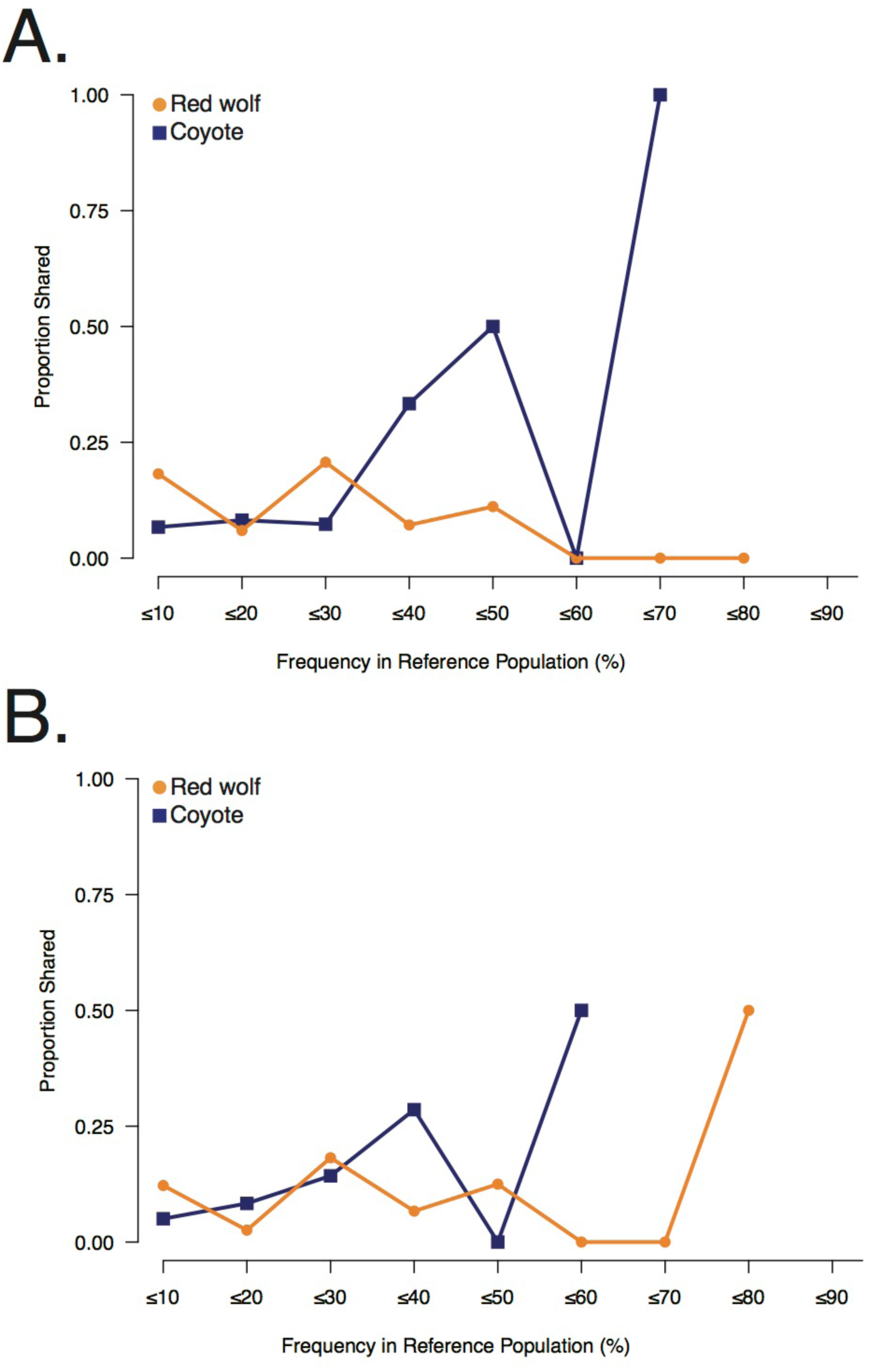
Sharing of red wolf and coyote private alleles with the two Galveston Island, Texas canids, A) GI-1 and B) GI-2, and their frequencies in their respective reference populations.

## Results

### Genome-wide SNP genotyping and diversity estimates

We collected genomic and mitochondrial (mtDNA) sequence data for two canids inhabiting Galveston Island, Texas (GI) of unknown taxonomic origin and 60 reference North American canids: 29 coyotes from the American southeast (Alabama, Louisiana, Oklahoma, Texas), 10 gray wolves from Yellowstone National Park, 10 eastern wolves (*C. lycaon*) from Algonquin Provincial Park in Ontario, and 11 red wolves from the Special Survival Plan captive breeding program (hereafter, red wolves) that collectively represent all extant red wolf founders (Table S1). We used a modified restriction-associated DNA sequencing (RADseq) protocol (*20*) to discover 7,068 genome-wide polymorphic SNPs after filtering for a minimum of three reads per allele, removing sites in high statistical linkage disequilibrium, sites with a minor allele frequency of 1%, and excluded sites with more than 10 % missing data.

We observed a significant difference in heterozygosity across all pairwise combinations of reference groups, where coyotes exhibited the highest genomic diversity of all reference groups (HE: coyotes=0.101, gray wolf=0.076, eastern wolf=0.087, red wolf=0.061; Tables 1, S1). In contrast, red wolves displayed the lowest levels of genomic diversity (H_E_=0.061), a trend consistent with expected erosion of diversity due to a declining effective population size, a subsequent bottleneck, and inbreeding (*21*). We found the highest levels of genomic differentiation between red and gray wolves (F_ST_=0.136), with the lowest levels found between coyotes and wolf comparisons (F_ST_: red wolf-eastern wolf=0.093, gray wolf-eastern wolf=0.086, coyote-gray wolf=0.062, coyote-red wolf=0.040, coyote-eastern wolf=0.042).

**Table 1.**
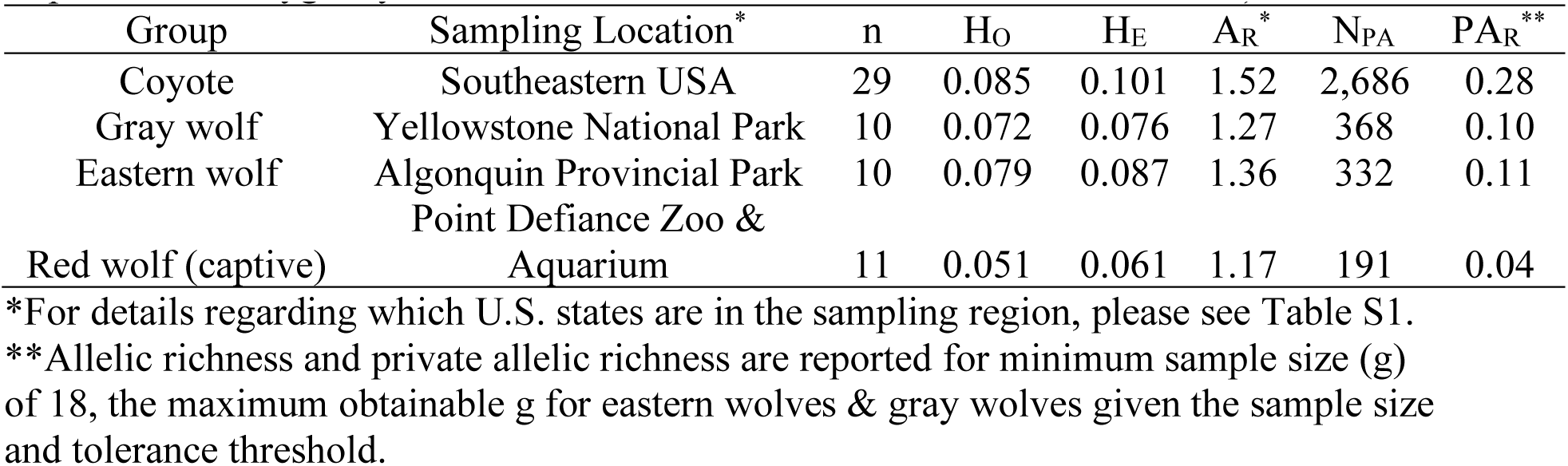
Diversity statistics for each reference group of *Canis* in North America. Summary statistics for each *Canis* reference population derived from 7,047 genome-wide polymorphic SNP_s_. (Abbreviations: n, sample size; HO, observed heterozygosity; H_E_, Expected heterozygosity; A_R_, Allelic Richness; PA_R_, Private Allelic Richness)

### Genetic clusters and assignment analysis

An initial principal component analysis (PCA) of all canids revealed that spatial clusters were largely concordant with taxonomic classifications (Fig. 1A) and consistent with previous genome-wide analyses of wolf-like canids (*22*). Specifically, PC1 explained 7.0% of the total variation, polarized by gray wolves and coyotes, with eastern and red wolves spatially intermediate, and the two GI canids proximal to the coyote cluster. The second axis was informative for red wolves (PC2, 3.9% variation), a pattern previously observed and likely attributed to the captive breeding bottleneck, inbreeding, and subsequent drift. When we restricted our analysis to only reference red wolves, reference coyotes, and the two GI canids, we found that coyotes and red wolves again defined PC1 (6.0% variation) with an intermediate placement of the two GI canids, and PC2 (4.4% variation) reflected geographic variation within coyotes (Fig. S1A).

We used a maximum likelihood framework in ADMIXTURE (*23*) to assess genetic structure across all 62 canids and found the greatest support for three genetic clusters (*K*=3, *cv*=0.35) composed of gray and eastern wolves, coyotes, and red wolves, respectively (Figs. 1C, S1B). The two GI canids exhibited partial memberships only to the red wolf and coyote clusters at *K*=3 (canid GI-1: *Q*_Red Wolf_=0.60, *Q*_Coyote_=0.40; canid GI-2: *Q*_Red Wolf_=0.60, *Q*_Coyote_=0.40). Interestingly, two coyotes that were collected in Louisiana also exhibited non-trivial assignment proportions to the red wolf genetic cluster (*Q*_Red Wolf_: LA-2=0.10; LA-3=0.11). A similar analysis of the GI canids with reference coyotes and red wolves showed strong support for two genetic clusters, which mirrored the patterns revealed by PCA (Fig. S1). Cluster memberships were similar as before, where clusters corresponded to a red wolf and coyote group, with the two GI canids assigned to each cluster (Fig. S1C). When we surveyed higher levels of partitioning, *K*=3 revealed two distinct groups of coyotes corresponding to their historic range of Oklahoma and Texas and to their southeastern expansion front across Louisiana and Alabama. At this level of partitioning, the two GI canids retained non-trivial assignments to red wolves (*K*=3 *Q*_Red Wolf_: GI-1=0.27, GI-2=0.21) (Fig. S2). Interestingly, the proportion of coyote ancestry for the two GI canids was attributed to the recently expanded southeastern coyote population (*Q*_Southeast Coyote_: GI-1=0.73; GI-2=0.79). Although no definitive conclusion can be drawn from this data regarding the origin on the red wolf allele sharing among the two GI canids, assignments to the southeastern coyote cluster, rather than the historic range cluster, is consistent with interbreeding between the wild red wolf and expanding coyote populations in the late 1970s.

We obtained the probability that each GI canid shared ancestry with reference coyotes, red wolves, or both using the posterior probability assignment test in STRUCTURE (*24*), where each GI canid was explicitly assigned to one or more of the coyote and red wolf reference groups. We found similar results, where each GI canid had partial assignments to the red wolf cluster (*Q*_Red_ Wolf: GI-1=0.33, GI-2=0.28) (Fig. S1C).

### Shared private alleles

We explored the degree to which each of the GI canids shared private alleles with the reference canid groups. This method is an informative approach to infer population relationships (*25*), where an excess of shared private alleles may imply source population or recent introgression (*26*). We surveyed 6,859 loci with non-missing data for canid GI-1 and estimated the proportion of alleles carried that are private to each reference group (Table 2). GI-1 shared the most private alleles with coyotes (S_PA_ n=184), followed by red wolves (S_PA_ n=21), with few shared private alleles with either gray wolves (S_PA_ n=12) or eastern wolves (S_PA_ n=12) (Table 2). This trend persisted following correction for unequal sample size (Coyote S_PAr_=0.0102; S_PAr_=0.0059) (Table 2). There was minimal sharing with other reference canid lineages (SPA_r_: *C. lupus*=0.0035; *C. lycaon*=0.0045). We surveyed 6,391 loci with non-missing data for canid GI-2 and found similar trends as for GI-1, where after correcting for sample size, again the greatest private allele sharing observed was between coyotes (S_PA_ n=138; S_PAr_=0.0093) followed by red wolves (SPA=14; S_PAr_=0.0063) and the other reference canid lineages (*C. lupus*: S_PA_ n=10; S_PAr_=0.0036; *C. lycaon*: S_PA_ n=8; S_PAr_=0.0039) (Table 2).

**Table 2.** Private allele sharing between reference groups and each Galveston Island, TX canid. Summary statistics for GI-1 and GI-2 were calculated over 6,859 and 6,391 genome-wide SNP_s_, respectively, reflecting the number of loci with non-missing data for each individual. (Abbreviations: N_PA_, number of private alleles; percentage; S_PA_, shared private alleles; S_Par_, shared private allelic richness)

| Reference Group | GI-1 |  |  |  | GI-2 |  |  |  |
| --- | --- | --- | --- | --- | --- | --- | --- | --- |
| | $N_{PA}$ | $S_{PA}$<br>(count) | $S_{PA}$<br>(%) | $S_{PAr}$ | $N_{PA}$ | $S_{PA}$<br>(count) | $S_{PA}$<br>(%) | $S_{PAr}$ |
| Coyote | 2,632 | 184 | 6.99% | 0.0102 | 2,439 | 138 | 5.66% | 0.0093 |
| Gray wolf | 362 | 12 | 3.31% | 0.0035 | 335 | 10 | 2.99% | 0.0036 |
| Eastern wolf | 329 | 12 | 3.65% | 0.0045 | 303 | 8 | 2.64% | 0.0039 |
| Red wolf | 188 | 21 | 11.17% | 0.0059 | 171 | 14 | 8.19% | 0.0063 |

As expected, the greatest proportion of shared private alleles between GI-1 and coyotes was observed for alleles with a frequency in coyotes of ≤70% (Proportion shared, prop_S_=1.00) (Fig. 2), and a low sharing ratio was observed for private alleles with low frequency in coyotes (e.g. ≤10%; propS=0.07) (Fig. 2). That is, GI-1 tended to share private alleles with coyotes that were relatively common across the coyote reference population. Interestingly, the same trend was not observed for private allele sharing with red wolves, where a relatively high proportion of allele sharing was observed for private alleles with ≤10% frequency in the reference population (prop_S_=0.18) (Fig. 2). However, the greatest proportion of allele sharing between red wolves and GI-1 was observed for private alleles at ≤30% frequency (prop_S_=0.21).

Similar trends were observed for GI-2, where the highest private allele sharing proportion with coyotes was observed for private alleles with ≤60% frequency (prop_S_=0.50) (Fig. 2) and the lowest proportion shared for alleles with ≤50% and ≤10% in the reference coyotes (prop_S_=0.00; 0.05) (Fig. 2). Additionally, when compared to red wolves, GI-2 had a moderately high private allele sharing proportion with low frequency private alleles (≤10%; prop_S_=0.12) (Fig. 2). However, the highest sharing proportion for GI-2 and red wolves was observed for private alleles with ≤80% frequency in the reference population (prop_S_=0.50) (Fig. 2).

As the GI canids predominantly shared private alleles with low to moderate frequencies in the red wolf population, this likely reflects the loss of diversity in red wolves due to the founding of the captive breeding population or subsequent inbreeding (*21,27*). Founders of the captive breeding population likely did not represent all red wolf diversity that existed on the landscape prior to the 1970s, when trapping to remove wild red wolves and founder selection occurred.

We observed 30 shared private alleles between at least one GI canid and red wolves. These markers were predominantly found in intergenic regions distributed across 21 chromosomes (n_intergenic_=19; n_exon_=3; n_intron_=7; n_promoter_=1) (Table S3). GI-1 was heterozygous at 76% of the loci that contained red wolf private alleles (n=16 out of 21 total sites), whereas GI-2 was heterozygous at 71% of the loci (n=10 of 14 total sites) (Table S3). We found five overlapping shared red wolf private alleles between the two GI individuals; however, we estimated a high level of allele sharing (IBS=0.93), likely due to originating from the same population and high probability of being related.

When the LD filter was removed, we retained 8,167 and 7,609 SNPs in GI-1 and GI-2, respectively. GI-1 carried a total of 30 red wolf private alleles (n_homozygous_=8) and GI-2 carried 26 (n_homozygous_=8) (Fig. S3). Although this provided a genome-level perspective of red wolf allele sharing, the resolution was not sufficient to conclusively identify contiguous shared private alleles in extended linkage disequilibrium due to recent admixture.

Overall, the GI canids carried 21 alleles that were absent from all reference populations. These GI canid-specific alleles were distributed throughout the genome (n, intergenic=16; exon=1; intron=4; promoter=0) (Fig. S3, Table S4B). GI-1 carried 14 such private alleles and was homozygous for 50% of loci, where GI-2 carried 16 private alleles and was homozygous for 56%. There were nine overlapping private alleles between the two GI canids, which again likely reflect a high probability of relatedness (Table S4B). It is possible that these alleles represent at least some of the genomic diversity in the historical wild red wolf population that was lost as the result of selecting 14 captive breeding founders from the wild, but this is speculation in the absence of documented historical red wolf samples.

### Matrilineal haplotyping

Canid mtDNA haplotypes are well established and form two clear gene tree clades: a clade composed of Eurasian-evolved gray wolves and domestic dogs, and a clade consisting of North American canids (coyotes, eastern wolves, and red wolves; Fig. 1D) (*28,29*). To further understand admixture in the GI canids and possible parental mtDNA contributions, we amplified the mtDNA control region that was previously found to contain a unique red wolf haplotype in contemporary wolves (*30*) and unique haplotypes in ancient canids of the American southeast (*31*). Both GI canids carried mtDNA haplotypes identical to previously published coyote haplotypes from the Great Plains states (GI-1: haplotype la77; accession JN982588; *32*) and Texas (GI-2: la143; accession FM209386; *17*). We reconstructed the two well supported mtDNA gene tree clades that are commonly identified with a high posterior probability (Prob=0.98) and found that the two GI canids clearly grouped with North American canids, specifically coyotes, although nodal support within the two clades was generally low (Figs. 1D, S4). For instance, the posterior probability was low (Prob<0.5) for most nodes within the North American clade, especially within coyote haplotypes, which show very little phylogenetic structuring across their range (*32*) (Figs. 1D, S4).

## Discussion

We rediscovered ghost red wolf alleles present on the Gulf Coast landscape 40 years after red wolves were believed to be extinct in the region. Through interbreeding with coyotes, endangered and extinct red wolf genetic variation has persisted and could represent a reservoir of previously lost red wolf ancestry. This unprecedented discovery opens new avenues for contemporary red wolf conservation and management, where ghost alleles could be re-introduced into the current captive and experimental populations. These admixed individuals consequently can be of great conservation value but there is no ESA policy providing protection for admixed individuals that serve as reservoirs for extinct genetic variation. An ‘intercross policy’ was introduced in 1996 to assist prioritizing protection efforts but this policy was never fully adopted (*33*). Several commentaries have encouraged an updated implementation of the ESA and Species Status Assessments, especially as admixed genomes are increasingly being described and viewed as a source of potentially beneficial genetic variation in the face of rapid climate change (e.g. *34*). Although red wolves represent one of the greatest species recovery stories in ESA history, debates regarding historical and on-going interbreeding with coyotes highlight the ESA’s short-comings associated with admixed individuals and the difficulty in setting management objectives given our evolving understanding of admixed genomes across wild populations (*35*).

In this study, our initial impetus to evaluate admixed individuals was due to the phenotype displayed by canids on Galveston Island, which appear to overlap with the canonical red wolf (Fig. 2). Given these suspected admixed canids, roadkill individuals were sampled for genomic testing. Our survey of their genomes revealed a surprising amount of allele sharing with the captive breeding population of red wolves. This shared variation could be the consequence of two potential scenarios: 1) surviving ancestral polymorphisms from the shared common ancestor of coyotes and red wolves that have drifted to a high frequency in the captive breeding red wolf population and in a small portion of Gulf Coast coyotes; or 2) wild coyotes in the Gulf Coast region are a reservoir of red wolf ghost alleles that have persisted into the 21^st^ century. Neither of these potential explanations require adherence to a specific species concept. For instance, incomplete lineage sorting from a shared common ancestor could occur whether red wolves are a subspecies of the gray wolf, conspecific with Eastern wolves, or an independent lineage with a possible ancient hybrid origin (*36*) (Fig. S5). Similarly, interbreeding with the ancestral red wolf population would have resulted in the introgression of red wolf alleles and associated phenotypes into Gulf Coast coyotes under each species concept. Our findings of admixture and composition of private alleles are most consistent with the second scenario, where the Galveston Island canids are admixed coyotes carrying red wolves ghost alleles. Further, Galveston Island is found within the historic red wolf range from where the original founders for the captive and reintroduced populations were captured in the 1970s (Fig. S6). On an island, this canid population likely experienced long-term reductions in gene flow with the southeastern coyote population and a greater probability of retaining unique red wolf alleles. In further support that coyotes of the American Gulf Coast likely serve as a ghost allele reservoir of red wolf ancestry, we also identified two coyotes with red wolf admixture from Louisiana’s Gulf Coast, a second geographic region in which trapping efforts were conducted to build a captive red wolf population (*16*). These findings provide substantial support that ancestral red wolf genetic variation persists as ghost alleles in the regional coyotes of the southeastern United States.

Our discovery warrants further genetic surveys of coyote populations in Louisiana and Texas to establish the level and extent to which remnant red wolf alleles are found exclusively in admixed coyotes. There are potentially admixed coyotes in the region that exhibit higher levels of red wolf ancestry. With these surveys in place, conservation efforts then face the opportunity to consider the role of remnant genetic variation in the future of the red wolf. The NCEP of red wolves are a listable entity under the ESA in need of proactive conservation (*36*). However, in the age of an extinction crisis, innovative mechanisms to preserve and utilize adaptive potential are in great demand. Today, every federally recognized red wolf individual is a descendant from 14 founders, of which only 12 are genetically represented. These founders were removed from a single geographic location in the 1970s and vastly underrepresent the original genomic diversity present in southeastern wolves (*5*). Our discovery of red wolf ghost alleles in southeastern coyotes demonstrate the ability to uncover ancestral variation and establish a new component of biodiversity conservation. A minority of conservation priorities have considered a ‘de-introgression’ strategy in which admixed individuals are bred in a specific design to recover the extinct genotype (*37*).

The “tree of life” approach to conservation is under challenge, as a new paradigm has been proposed to include admixed genomes (*35,38*). Historical and contemporary red wolves face anthropogenically-mediated hybridization, but introgression is also likely a natural process in the evolution of *Canis* lineages. As an important evolutionary process, introgression could protect adaptive potential and maintain processes that sustain ecosystems. As a result, incorporating admixed entities into conservation policy and here, red wolf restoration, may be the next step in broader biodiversity conservation. Another pivotal next step in red wolf restoration is the identification of a new re-introduction site for a wild population of red wolves. Our discovery of red wolf ghost alleles indicates there are geographic regions that can harbor endangered genetic variation and may guide future efforts for red wolf reintroduction. The foundation upon which that effort will be built rests exclusively on describing large-scale geographic patterns of red wolf ghost alleles in the American southeast.

## Materials and Methods

### Genomic DNA preparation and sample selection

We obtained tissue samples from two roadkill canids of unknown taxonomic affiliation on Galveston Island, TX (GI canids hereafter). We extracted genomic DNA using a DNeasy Blood and Tissue Kit (Qiagen) following the manufacturer’s instructions. We selected reference samples that collectively represented all possible wild canid evolutionary lineages in North America that could have contributed to the ancestry and genetic variation of the two GI canids (Table S1). The reference tissue samples were obtained via helicopter darting and post-mortality sampling by the US National Park Service (*C. lupus*), through the Ontario Ministry of Natural Resources archives (*C. lycaon*), or the Point Defiance Zoo and Aquarium archives (*C. rufus*). Reference coyote samples were obtained through the Sam Noble Oklahoma Museum of Natural History, samples collected from animal captured for radio collaring, or donations from hunters and trappers. We selected captive red wolves that captured genetic diversity of the 12 original founders and who are genetically represented in extant wolves that reproduced in the captive breeding program. These captive red wolves are considered the reference red wolves in all analyses.

### RADseq library preparation and bioinformatic processing of sequence data

We used a modified version of the RADseq protocol by (*20*). We digested DNA with *Sbf1*, ligated a unique barcode adapter to the fragments, and pooled between 96 and 153 samples. Each pool was subsequently sheared to 400 bp in a Covaris LE220 at Princeton University’s Lewis Siegler Institute Genomics Core Facility. We recovered ligated fragments using a streptavidin bead binding assay and prepared genomic libraries for Illumina HiSeq sequencing following either the standard TruSeq protocol for the NEBNext Ultra or NEBNext UltraII DNA Library Prep Kit (New England Biolabs). We conducted a size selection step using Agencourt AMPure XP beads (Beckman Coulter) to retain fragments 300-400 bp in size. We also used AMPure XP beads for library purification. We standardized genomic libraries to 10nM for 2X150nt sequencing on an Illumina HiSeq 2500 platform.

We filtered raw paired-end sequence data to retain reads that contained one of the 96 possible barcodes and the expected restriction enzyme cut-site using a custom perl script (flip_trim_sbfI_170601.pl, see Supporting Information). We discovered variant sites following the STACKS v 1.42 pipeline (*39*). Reads were de-multiplexed using *Process_Radtags*, allowing a mismatch of two to rescue barcodes. We discarded reads with an uncalled base or with an average quality score (≤10) within a sliding window equivalent to 15% of the total read length. We removed PCR duplicates using *Clone_Filter* with default parameters. All samples were mapped to the Canfam3.1 assembly of dog genome (*40*) with STAMPY v 1.0.31 (*41*). We filtered mapped reads in Samtools v 0.1.18 (*42*) to retain those with MAPQ >96 and exported as a bam file. Variant calling was completed in STACKS following a standard pipeline for reference mapped data (i.e. *pstacks*, *cstacks*, *sstacks*, *populations*) (*39*). We required a minimum stack depth of 3 reads (-m) in *pstacks* and allowed a maximum per locus missingness of 10% in *populations*. Further, to reduce biases resulting from linked markers, we enabled the *–write-single-snp* flag in *populations* and filtered for statistical linkage disequilibrium (LD) across sites using the *--indep-pairwise* 50 5 0.5 flag in Plink v1.90b3i (*43*). We conducted a final filtering to retain sites that also had a minimum minor allele of 1%.

Standard metrics of genomic diversity (observed heterozygosity, HO; private allele count, N_PA_, Pairwise F_ST_) for all reference groups were calculated using functions in STACKS. We evaluated significant differences in genome-wide heterozygosity pairwise estimates with a pairwise Wilcoxon rank sum test implemented in *R* with a false discovery rate correction for multiple testing (FDR<0.05). Allelic richness (A_r_) and private allelic richness (PA_r_) were calculated using a rarefaction approach implemented in ADZE (*25*) with a maximum tolerance of 10% missing data and maximum standardized sample size (g) set to the smallest n for the samples considered (20; eastern and gray wolves).

### Mitochondrial DNA sequencing and haplotyping

Mitochondrial sequence was amplified from genomic DNA with primers for control region (Thr-L 15926: 5′-CAATTCCCCGGTCTTGTAAACC-3′; DL-H 16340: 5′-CCTGAAGTAGGAACCAGATG-3) and thermocycling conditions following (*28*). Amplified products were bi-directionally sequenced using a service provided by GeneWiz (New Jersey). Each sample was sequenced in duplicate to confirm ambiguous sites. Sequences were view, corrected, and aligned with Geneious v6.16 software (*44*).

### Population Structure Analyses

To visualize genetic clustering patterns of the filtered SNP dataset, we conducted a Principal Component Analysis (PCA) in *flashPCA* (*45*). Additional PCAs were completed on a subset of the total sample size for specific comparisons (i.e. inclusion of only reference coyotes, reference red wolves, and the two GI individuals). We implemented a maximum-likelihood analysis to infer population structure using the program ADMIXTURE v1.3. (*23*). We evaluated between 1 and 10 genetic partitions (*K*), evaluated the fit of each partition using the cross-validation flag, and considered the best fit number of partitions to have the lowest cross-validation score. We first considered the entire dataset, with subsequent analyses conducted on subsets of the total sample size for specific comparisons. For example, we analyzed only the reference coyotes, reference red wolves, and the two GI canids using the aforementioned parameters.

Although this maximum-likelihood cluster analysis is useful for evaluating specific levels of data partitioning, it is not an explicit ancestry analysis. Using a Bayesian framework, we conducted a posterior probability assignment test in Structure v. 2.3.4 (*24*) that included all reference coyotes and red wolf individuals as the training set of samples. We then assigned each of the GI canids to one or more of these reference groups (*K*=2) using 10,000 repetitions following a burn-in of 2,500.

### Private Allele Sharing Analyses

We evaluated the degree of sharing of private alleles among the GI canids and all possible reference groups, considering each GI canid separately. To avoid spurious identification of private alleles due to the presence of any missing data, we restricted analyses to loci that were 100% genotyped in each GI canid and identified alleles private to each reference group in STACKS. We then determined the number of shared private alleles between the GI canid and the reference groups. Further, we calculated shared private allelic richness with each reference group using a rarefaction approach in ADZE with a tolerance of 15% missing data and a maximum standardized sample size (g) of two.

We estimated the frequency of each shared private allele in the corresponding reference coyote or red wolf population. This frequency distribution was binned in frequency intervals of 10%. In other words, the number of shared private alleles for each GI canid was divided by the total number of private alleles and binned in 10% frequency intervals based on the allele’s frequency in the corresponding reference population.

We determined the genomic coordinates of all shared red wolf private alleles with each of the GI canids. We annotated each site as intergenic, within an intron or exon, or within a putatively regulatory region (within 2 kb of a transcription start site) using a custom python script and the Ensembl gene database (chr_site.py; see Supporting Information) (*46*). To further evaluate the fine scale distribution of shared red wolf private alleles across the genomes of GI-1 and GI-2, we removed the LD filter and recalculated shared private alleles with the red wolf reference group as described above. We evaluated the alleles found only in GI-1 and GI-2 to determine whether either GI canid harbored any unique genomic diversity absent from the reference groups. We calculated the identity by state (IBS) between the two GI canids in Plink v1.90b3i using the --ibs-matrix argument.

### Mitochondrial DNA sequence analysis

We compared the GI canids mtDNA haplotypes to mtDNA control region sequences available on GenBank the represented possible *Canis* ancestors from multiple evolutionary lineages: domestic dogs, gray wolves, eastern wolves, red wolves, and coyotes (Table S2). Using consensus sequences of 234bp, we estimated gene trees using Bayesian methods implemented in BEAST v1.8.4 with red fox (*Vulpes Vulpes*) as an outgroup (*47*). In BEAST, we use a constant size coalescent tree prior, an uncorrelated lognormal relaxed molecular clock, and used a random starting tree. We conducted two independent Markov Chain Monte Carlo (MCMC) analyses for 25 million steps, sampling every 2,500 steps, and combined tree estimates from each run with LogCombiner v1.8.4 with a 10% burnin. Convergence on the posterior distribution was determined based on viewing the log files in Tracerv1.6, where convergence is attained when the effective sample size of a parameter (i.e. the number of effectively independent draws from the posterior distribution) is at least 200; all parameters in our analyses had effective sample size values greater than 400. To visualize our estimated gene trees, we first calculated the maximum clade credibility in TreeAnnotator v1.8.4 and upload the most likely tree in the Interactive Tree of Life v3.6.3 online platform (*48*). Full tree results are presented in Figure S5 with posterior probabilities over 0.90 reported; truncated results that show the clearly detected phylogenetic relationships among North American canids and GI canids are presented in Figure 2D.

## Acknowledgments

We are thankful to Oklahoma Museum of Natural History, who provided SE coyote samples under loan 501 agreement G.2016.3, 4318. We also want to thank Brent Patterson for the use of eastern wolf samples in this study. We also thank L. David Mech, Ronald M. Nowak, and Cornelia Hutt for assistance with and discussion about the initial discovery of Galveston Island canids. Funding provided by The Point Defiance Zoo and Aquarium Dr. Holly Reed Conservation Fund, the National Science Foundation Postdoctoral Research Fellowship in Biology under Grant No. 1523859, and the Ontario Ministry of Natural Resources and Forestry, Natural Sciences and Engineering Research Council of Canada. DRS was funded by NSF (DEB-1245373) and Yellowstone Forever.

## Data and materials availability

All data needed to evaluate the conclusions in this paper can be obtained from the National Center for Biotechnology Information Sequence Read Archive (SUB4398163). Additional data can be requested from the corresponding authors.

